# Machine Learning Unravels Inherent Structural Patterns in *Escherichia coli* Hi-C Matrices and Predicts DNA Dynamics

**DOI:** 10.1101/2023.12.20.572497

**Authors:** Palash Bera, Jagannath Mondal

## Abstract

The large dimension of the Hi-C-derived chromosomal contact map, even for a bacterial cell, presents challenges in extracting meaningful information related to its complex organization. Here we first demonstrate that a machine-learnt (ML) low-dimensional embedding of a recently reported Hi-C interaction map of archetypal bacteria *E. Coli* can decode crucial underlying structural pattern. In particular, a three-dimensional latent space representation of (928*×*928) dimensional Hi-C map, derived from an unsupervised artificial neural network, automatically detects a set of spatially distinct domains that show close correspondences with six macro-domains (MDs) that were earlier proposed across *E. Coli* genome via recombination assay-based experiments. Subsequently, we develop a supervised random-forest regression model by machine-learning intricate relationship between large array of Hi-C-derived chromosomal contact probabilities and diffusive dynamics of each individual chromosomal gene. The resultant ML model dictates that a minimal subset of important chromosomal contact pairs (only 30 %) out of full Hi-C map is sufficient for optimal reconstruction of the heterogenous, coordinate-dependent sub-diffusive motions of chromosomal loci. Specifically the Ori MD was predicted to exhibit most substantial contribution in chromosomal dynamics among all MDs. Finally, the ML models, trained on wild-type *E. Coli* was tested for its predictive capabilities on mutant bacterial strains, shedding light on the structural and dynamic nuances of ΔMatP30MM and ΔMukBEF22MM chromosomes. Overall our results illuminate the power of ML techniques in unraveling the complex relationship between structure and dynamics of bacterial chromosomal loci, promising meaningful connections between our ML-derived insights and real-world biological phenomena.

## I. INTRODUCTION

The archetypal bacterium *Escherichia coli* (*E. coli*) possesses a super-coiled circular DNA with a length of 1.6*mm* and a size of 4.64 Mega basepair (Mb), confined within a (24)*μm* long spherocylinder [1, 2]. Over the years, our understanding of the *E. coli* chromosome has evolved significantly. Initially it was thought that chromosome is a just like a complex blob of various macromolecules such as DNA, proteins, RNA, etc. However, subsequent findings [3–6] reveal that, instead of a complex, blob-like architecture, it consists of a well-organized structure with distinct domains known as macro domains(MDs) [7–12]. In this regard, various chromosome conformation capture techniques [13–15] provide us with crucial information, unraveling the spatial organization of the genome, especially in understanding higher-order structures. Recent upgrade in high throughput genome sequencing technique (Hi-C) allows to investigate the three-dimensional conformation of *E. coli* chromosomal DNA [16]. This innovative method generates a high-resolution contact map, referred to as the Hi-C matrix which captures the proximity and frequency of contact between various regions of *E. coli* chromosome. Furthermore, the Hi-C matrices for different mutant provide valuable insights into the functions of nucleoid-associated proteins (NAPs) and their roles in maintaining the nucleoid’s structure [16]. The resulting chromosomal organization significantly influences the dynamic behaviour of chromosomal loci. Previous fluorescence based experimental studies have revealed that these chromosomal loci move subdiffusively [17–20], with their motion strongly influenced by genomic coordinates, showcasing a remarkable level of heterogeneity in their dynamics [19]. Integrating this Hi-C-derived contact information into a polymer-based model has enabled theoretical studies to furnish a plethora of structural and dynamical details regarding the *E. coli* chromosome [21–24]. These theoretical studies achieve a level of experimental accuracy that enhances our comprehension of the intricacies governing the organization and dynamical behavior of the *E. coli* chromosome.

The Hi-C-derived chromosomal contact map represents a multi-dimensional interaction matrix [25, 26]. Even for a prokaryotic cell such as *E. coli*,with singular circular DNA of 4.6 Mb sequence-length, the Hi-C matrix [16] manifests a dimension as large as (928*×*928) at a 5 Kb resolution. As a result, discerning meaningful information via its visual inspection of extremely large dimensional heterogenous interaction map can be challenging. In recent years, the state of the art machine learning (ML) techniques have emerged as powerful tools for automated extraction of valuable insights from large dimensional data. In light of this, we revisit the Hi-C derived contact probability map of *E. coli* and pose following questions :

- What underlies a ML-derived low-dimensional representation of the Hi-C map of *E. coli* chromosome?
- Can we quantitatively extract a subset of minimal chromosomal contact informations that would sufficiently reconstruct the experimentally observed [19] heterogeneous sub-diffusive motion of chromosomal loci?
- To what extent would the ML-based learning of wild-type chromosomal contact information aid in the prediction of Hi-C map of NAP-devoid mutant?

To address these questions, we first employ an artificial neural network (ANN) based frame-work in a bid to uncover crucial structural insights embedded within this large Hi-C matrix. As would be revealed in first part of Results section, a latent space representation of the Hi-C map successfully identified various MDs with a high degree of accuracy with experimentally derived MDs. In the later part of the manuscript, in a complementary approach, we integrate Hi-C contacts into a polymer-based model, predicting diffusive dynamics of a large number of chromosomal loci using a supervised machine learning technique called Random Forest (RF) regression. As would be unveiled in the manuscript, the proposed regression model successfully recover the coordinate-dependent heterogeneous subdiffusion [19] of chromosomal loci. Moreover, we extract important features from the input data that are crucial in maintaining this dynamical behavior of the loci. By incorporating only these important features related to Hi-C contacts into the polymer model, we successfully reproduce loci dynamics. Finally, we provide our insight on extent of predictive ability of both structure and dynamics of two NAP-devoid mutants of the chromosome (namely ΔMatP30MM and ΔMukBEF22MM) by ML models trained on wild-type chromosome.

## II. RESULTS

### A. Unsupervised ML model identifies a meaningful intrinsic structural pattern embedded within the Hi-C matrix of *E. Coli* chromosome

#### Overview of ML architecture

We employed an unsupervised machine learning algorithm known as Autoencoder [31, 32] to unveil the essential structural insights embedded within the Hi-C matrix. Autoencoder is a type of unsupervised deep neural network characterized by a dual structure comprising an encoder and a decoder, with a bottleneck in between (Figure 1(a)). In this architecture, the encoder converts the input data from a high-dimensional space to a lower-dimensional representation known as the latent space. Subsequently, the decoder reconstructs the initial input data from this latent space. This process involves the adjustment of model parameters, primarily weights and biases. Consequently, the compressed representation within the latent space reflects a non-linear transformation of the original input data, encapsulating crucial information or patterns inherent in the input datasets.

**FIG. 1.**
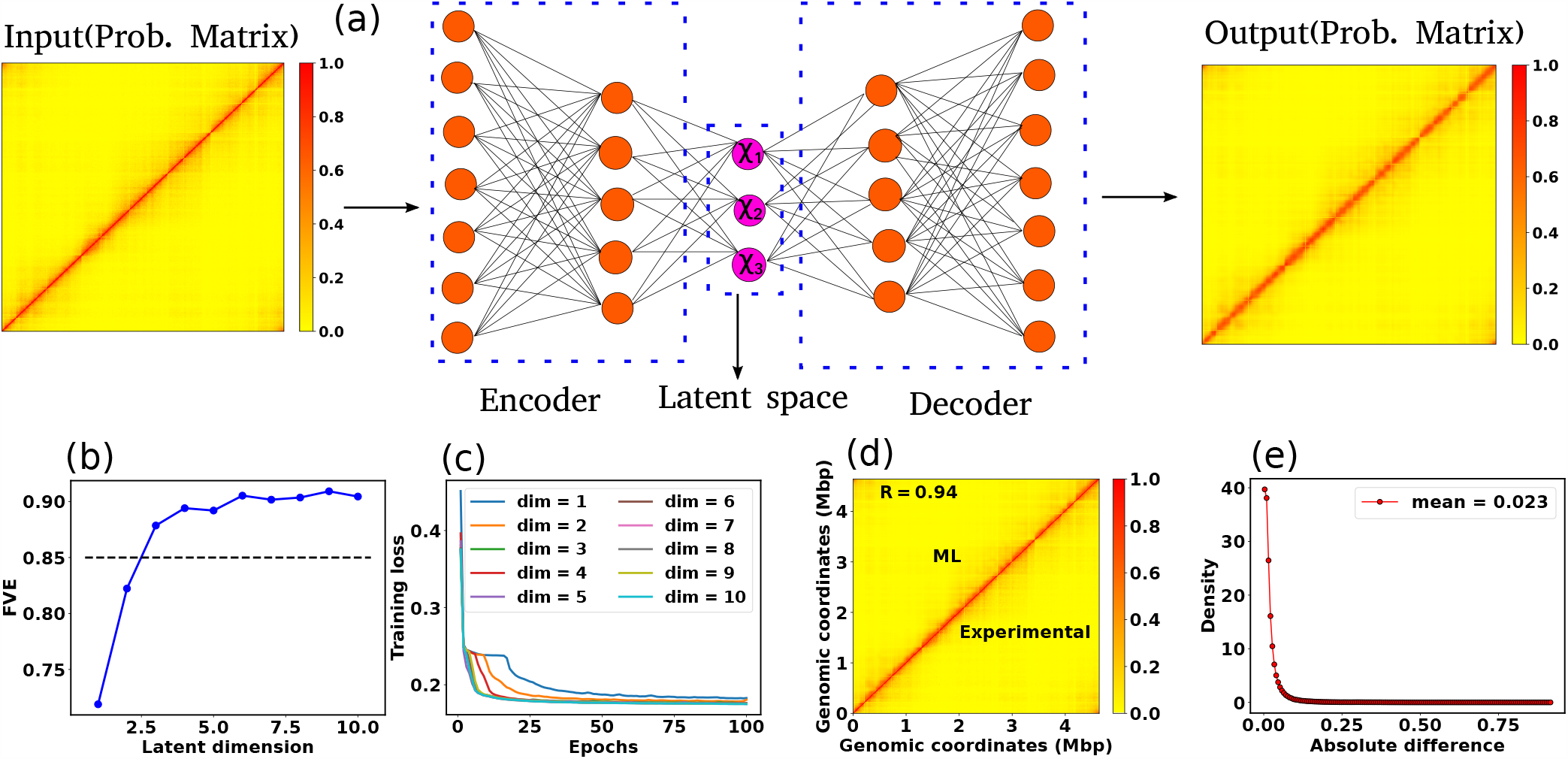
Architecture of the Autoencoder and training robustness. (a) Schematic of the Autoencoder, an unsupervised machine learning algorithm. It consists of an encoder and a decoder and in between there is a bottleneck. The encoder transforms high-dimensional input data to a lower-dimensional latent space, while the decoder reconstructs the initial input data from the latent space. This process involves adjusting model parameters, primarily weights, and biases. Each dimension in the latent space corresponds to a latent variable. Here *χ*_1_, *χ*_2_, and *χ*_3_ represent three latent variables. (b) The variation of FVE with respect to the latent dimension (*L*_*d*_). A *L*_*d*_ = 3 was chosen, ensuring an FVE of at least 0.85, signifying that the Autoencoder’s reconstruction captures a minimum of 85% of the variance in the input data. (c) Training loss as a function of epochs for different latent space dimensions (*L*_*d*_). Notably, the training loss achieves saturation for all *L*_*d*_ beyond 25 epochs. (d) Genome-wide contact probability map between the experimental and ML-derived Hi-C matrix. (e) Histogram of the absolute difference between experimental and ML-derived contact probability matrices. The Pearson correlation coefficient(PCC) is 0.94, and the absolute difference in mean values is 0.023, indicating a substantial agreement in chromosomal interactions.

In our ML model, the input comprises a single Hi-C probability matrix with dimensions 928*×*928 (4640kb/5kb = 928). The Autoencoder architecture is structured with a total of nine sequential layers featuring neuron counts of 928, 500, 200, 100, *L*_*d*_, 100, 200, 500, and 928, respectively, where *L*_*d*_ denotes the dimension of the latent space. After setting up the Autoencoder architecture, we need to choose *L*_*d*_ judiciously. To determine this dimension, we computed the Fraction of Variance Explained (FVE) through reconstruction, which is defined as

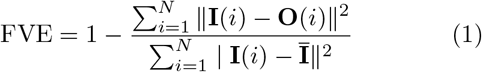

Here, **I**(*i*), **O**(*i*), and **Ī** represent the input, output, and mean input, respectively, and *N* = 928 corresponds to the number of rows in the Hi-C matrix. Figure 1(b) represents the variation of the FVE as a function of latent dimension. We opted for a latent dimension of *L*_*d*_ = 3, as it helps to achieve an FVE of at least 0.85, meaning that the Autoencoder’s reconstruction accounts for a minimum of 85% of the variance in the input Hi-C data. This choice of latent space dimension not only ensures effective data representation but also affords flexibility in visualizing the compressed data.

To assess the training robustness across various latent space dimensions (*L*_*d*_) concerning the number of epochs, we have plotted the training loss as a function of epochs for different *L*_*d*_ (Figure 1(c)). The figure clearly illustrates that beyond epochs = 25, the training loss reaches a point of saturation for all *L*_*d*_. This observation implies that selecting a number of epochs greater than 25 is a prudent choice. In our model, we opted for 100 epochs and in the *Method section*, additional specifics regarding the training of the Autoencoder are discussed. In a similar vein, we conducted a comparison between the input (experimental) and output (reconstructed) matrices. Figures 1(d) and (e) compare the genome-wide contact probability map between the experimental and ML (reconstructed by the Autoencoder with *L*_*d*_ = 3) contact probability matrix, along with a histogram showing the difference between the two matrices. Our findings reveal a Pearson correlation coefficient(PCC) of 0.94 between the experimental and ML contact probability matrices. Additionally, the absolute difference in the mean values is 0.023, indicating a substantively strong agreement between experimental and ML-derived chromosomal interactions.

#### The pattern emergent from latent space of the ML-model recovers key Macro-domains across *E. coli* genome

We aim to understand the biological significance of the lower-dimensional representation (*L*_*d*_ = 3) of the input data. To achieve this, we generated a scatter plot of the latent space data and conducted clustering using the K-means algorithm [33, 34]. Our hypothesis is that each cluster signifies specific domains within the bacterial chromosome, inherently encoded in the Hi-C matrix. Biologically, these large-scale structurally distinct domains are referred to as macrodomains(MDs) [7–12]. It is noteworthy that the actual molecular mechanisms governing macrodomain organization remain incompletely understood, and the precise boundaries of these MDs have been found to vary across different reports [7–11].

In Figure 2(a), a scatter plot illustrates the three dimensional (*L*_*d*_ = 3, *χ*_1_, *χ*_2_, and*χ*_3_) representation of the latent space, with distinct color-coded clusters representing various MDs of the chromosome. From experimental study [10], we possess a priori knowledge regarding the base pairs of each macrodomain. Additionally, through clustering, we have obtained base pair information for each macrodomain. Subsequently, we conducted a detailed comparison between experimentally denoted and (ML)-derived MDs by schematically drawing the DNA as a circle (Figure 2(b)). The inner and outer circles, featuring various color-coded regions, represent the experimentally denoted and ML-derived macrodomains, respectively, with base pair information annotated in kilo bases (kb). A visual inspection indicates substantial agreement between MDs, barring discrepancies in the NSR, Right, and Ter MDs. Quantitative comparison between actual (experimentally denoted) and predicted (ML-derived) MDs is facilitated by the confusion matrix (see SI for details). Metrics such as Accuracy, Precision, Recall, and F1-Score can be computed from the confusion matrix as follows.

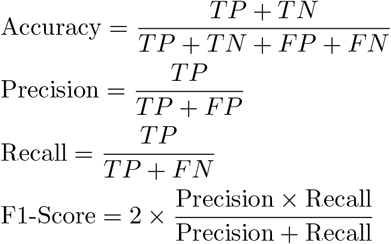

**FIG. 2.**
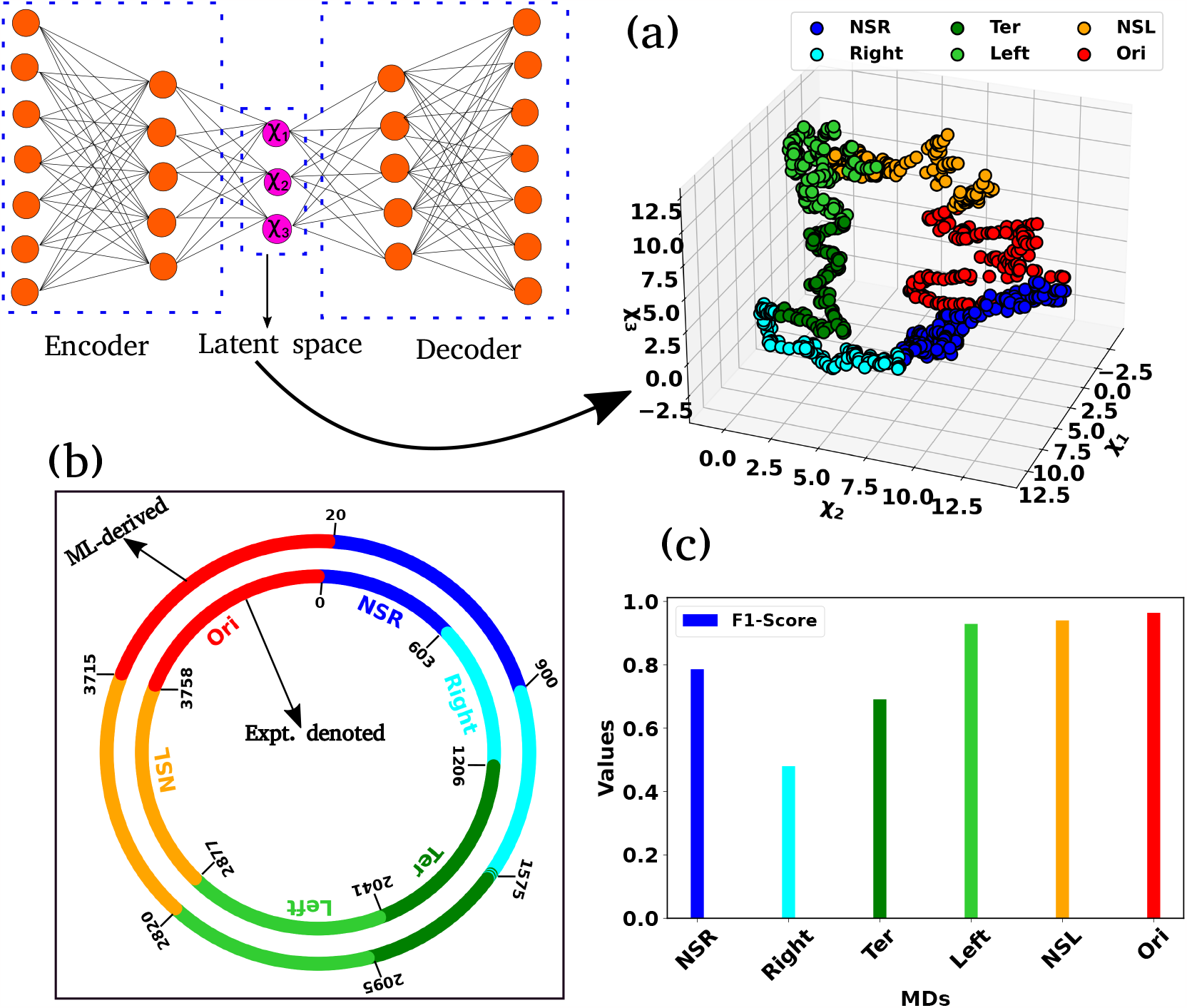
Representation of the latent space and classification of different macrodomains. (a) Three dimensional scatter plot of the latent space variable *χ*_1_, *χ*_2_, and *χ*_3_. The data has been clustered using K-means clustering. The various color-coded clusters are representing distinct macrodomains (MDs) within the bacterial chromosome. (b) The comparison between experimentally denoted and machine learning (ML)-derived MDs. The inner and outer circles, each encoded by various color-coded regions, delineate the experimentally denoted and ML-derived macrodomains, respectively, with base pair information annotated in kilo bases (kb). (c) The bar plot of the F1-score for different MDs. The higher values of the F1-Score indicated the better classification.

Figure 2(c) shows a bar plot of the F1-score for each MD. F1-scores exceeding 0.92 for the Left, NSL MDs suggest a strong match between actual and predicted classes. Conversely, lower F1-Scores for the other three MDs indicate a moderate alignment, consistent with observations in

Figure 2(b). Nevertheless, the overall accuracy for all classes stands at 0.82, indicative of a robust correlation between experimentally denoted and ML-predicted MDs. In summary, our unsupervised ML model offers a potent automated approach for MDs identification, demonstrating a high degree of accuracy with experimentally derived MDs.

### B. ML-based identification of genomic contacts crucial for *E. Coli* heterogeneous dynamics

In the preceding section, we delved into the intrinsic structural properties of *E*.*Coli* chromosome embedded within the Hi-C matrix, which led to an automated discovery of segmented macrodomains in a ML-derived low-dimensional subspace. In this section, we now pose the question: Can we identify the crucial subset of chromosomal contacts in Hi-C map, that hold key to the heterogenous, cooordinate-dependent diffusivities [19, 22] of chromosomal loci. Towards this end, we employed a ML-based protcol namely Random Forest Regression to extract dynamical information by leveraging the structural properties of the chromosome, such as the pairwise distance between chromosomal beads, This unsupervised, trees-based algorithm, initially proposed by Breiman et al. [35, 36], is a potent tool widely used for both classification and regression tasks. Random Forest Regression operates by randomly selecting input data from training datasets and creating an ensemble of trees (forests) based on these features and labels of the input data. These ensembles of trees are called “decision trees”. The final results are derived by averaging (for regression) or voting (for classification) from the outputs of these decision trees. Notably, the Random Forest possesses a distinctive ability to pinpoint the most crucial features within the training datasets.

#### Data preparation and supervised ML architecture

We implemented a bead-in-a-spring polymer model to simulate the bacterial chromosome and generated a set of 200 distinct initial DNA configurations. In brief, the resolution of each bead is 5*×*10^3^ bp (5 kb), similar to the Hi-C matrix resolution [16]. Each bead has a diameter denoted by *σ*, and the chromosome is confined within a spherocylindrical boundary that mimics the cell wall. The bonded interactions between adjacent beads have been modeled by harmonic springs, while the non-bonded interactions are represented by the repulsive component of the Lennard-Jones potential *V*_*nb*_(*r*) = 4*t:*(*σ/r*)^12^, where *t:* is the potential depth, and *r* is the distance between two beads. The Hi-C interactions are also modeled as effective springs with a spring constant and bond lengths dependent on the strength of the contact probabilities. Following energy minimization of the initial configurations, Brownian Dynamics simulations are conducted for each configuration at a temperature of *k*_*B*_*T* = 1.0 and friction *γ* = 1.0. The length and time scales are represented in the unit of *σ* (the diameter of each bead) and 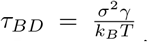 = (Brownian time), respectively throughout the manuscript. The simulations are run for a time duration of 10^3^*τ*_*BD*_, ensuring proper equilibration of each configuration. After equilibration, the pairwise distances are computed using the last snapshot of each run (totaling 200), serving as features for our machine-learning model. The dynamics of each DNA bead are quantified by calculating the mean squared displacement (MSD). For this measurement, we simulate each equilibrium configuration 40 times through Brownian dynamics simulations, drawing distinct velocities from the Maxwell-Boltzmann distribution at a desired temperature of *k*_*B*_*T* = 1.0. These trajectories, with varying initial velocities, are called isoconfigurational ensembles [37, 38]. We computed the MSD of each particle *i* and averaged over the different runs (isoconfigurational ensembles) i.e.

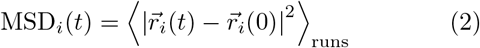

where 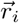 represents the position of *ith* particle and angular bracket signifies the average over isoconfigurational ensembles. So from the equilibrium configuration, we have calculated the pair-wise distance of the chromosomal beads and the MSDs of individual beads from the isoconfigurational ensembles. These two quantities serve as features and labels, respectively, for our machine-learning algorithm, Random Forest. By using this technique we can predict the dynamics of bacterial chromosomal loci and extract the important chromosomal contact features which are necessary to maintain the dynamics. The entire process of data preparation and the machine learning architecture are schematically depicted in Figure 3(a) and (b), respectively. In our ML model, we allocate 75% of the data for training and 25% for testing. This partitioning ensures that the dimensions of the training and testing data are as follows: features_training_[*m, n*] = [runs *×* 928, 928] = [150 *×* 928, 928], labels_training_[*m*] = [runs*×*928] = [150*×*928] and features_testing_[*m, n*] = [runs *×* 928, 928] = [50 *×* 928, 928], labels_testing_[*m*] = [runs *×* 928] = [50 *×* 928].

**FIG. 3.**
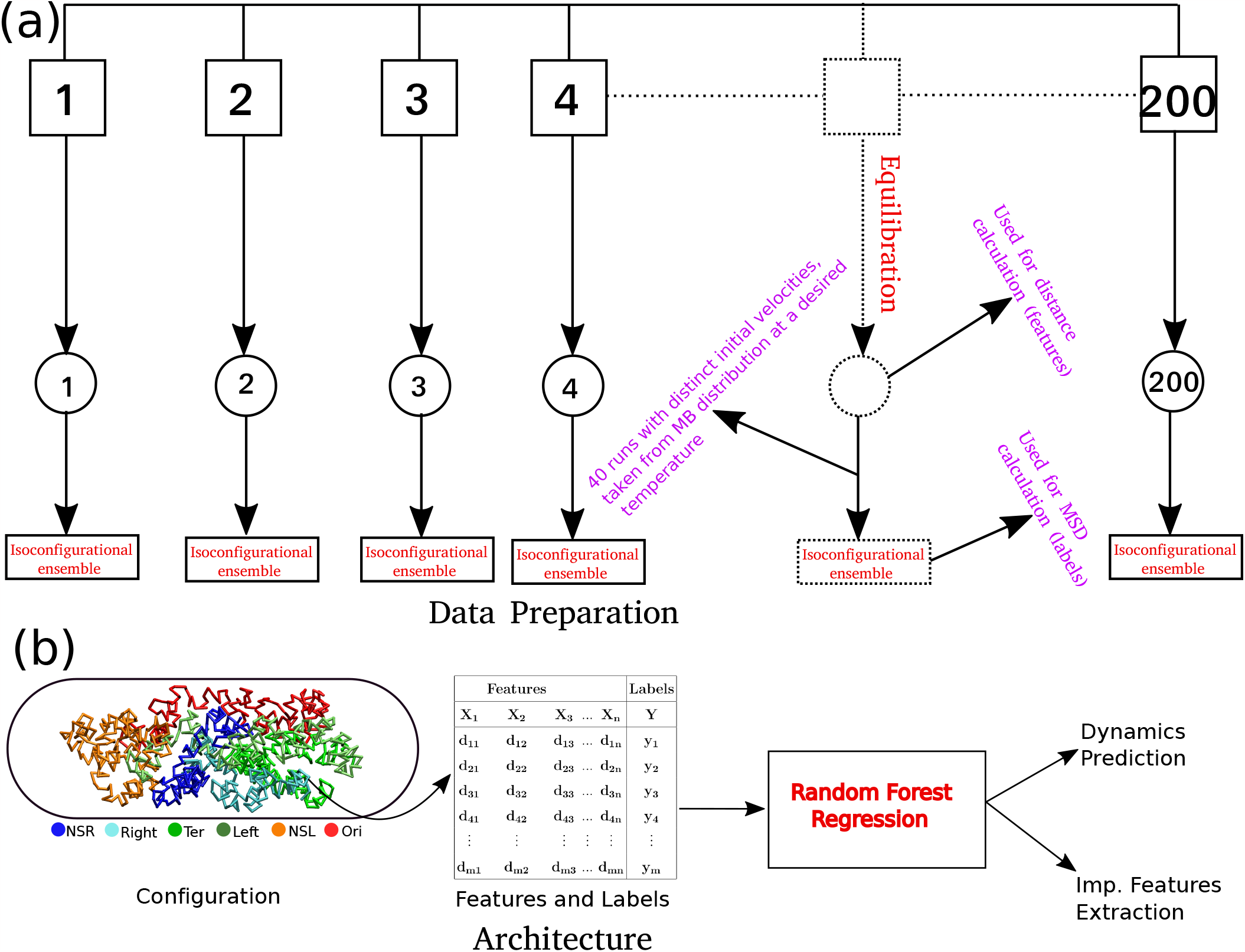
Schematic representation of the Data preparation and the architecture of Random Forest (RF) regression model. (a) Data Preparation: At the very beginning we generated a set of 200 distinct initial configuration of DNA using bead-in-a-spring polymer model. We run the Brownian dynamics simulation for each configuration for a time span of 10^3^*τ*_*BD*_, to ensure proper equilibration. Following equilibration, pairwise distances were computed using the last snapshot of each run (totaling 200), serving as features for our machine-learning model. For the MSD measurement, we simulate each equilibrium configuration 40 times through Brownian dynamics simulations, drawing distinct velocities from the MaxwellBoltz-mann distribution at a desired temperature of *k*_*B*_*T* = 1.0. These ensemble (40 trajectory each) are called iso-configurational ensembles. (b) Architecture: We utilized the pair wise distance between and MSDs of each beads. These two quantities serves as features and labels respectively, in our ML model. After training of the Random Forest (RF) with this datasets, we predicted the dynamics of individual beads. For training and testing, 75% and 25% of the total datasets, were used. Additionally, from the trained model, we extracted important features contributing to the maintenance of dynamical properties.

#### Comparison between the actual and ML-predicted MSDs of different loci and their exponents

Upon training the Random Forest regression model, we proceeded to predict the dynamics of individual chromosome loci. The selection of specific loci was based on a prior experimental study conducted by Javer et al. [19], which suggested that chromosomal loci belonging to the Ter region exhibit slower motion, while those in the Ori region demonstrate faster motion. To assess the accuracy of ML predictions, we computed the PCC, denoted by *ρ*, between the predicted and actual MSD values. Figure 4(a) shows the variation of *ρ* as a function of time. Remarkably, the correlation *ρ* exhibits higher values (*>* 0.8) for shorter time intervals. However, the correlation shows a slight decrease for longer duration. Now we will delve into the dynamic properties of distinct loci within the DNA. Each macro domain is comprised of various loci identified by Espeli et al. [10] based on their genomic coordinates. Figure 4(b) represents the comparison between the actual and predicted MSD as a function of time for two distinct loci, Ori2 and Ter3. This figure distinctly reveals a close alignment between the actual MSDs (solid line) and the predicted MSDs (dotted line).

**FIG. 4.**
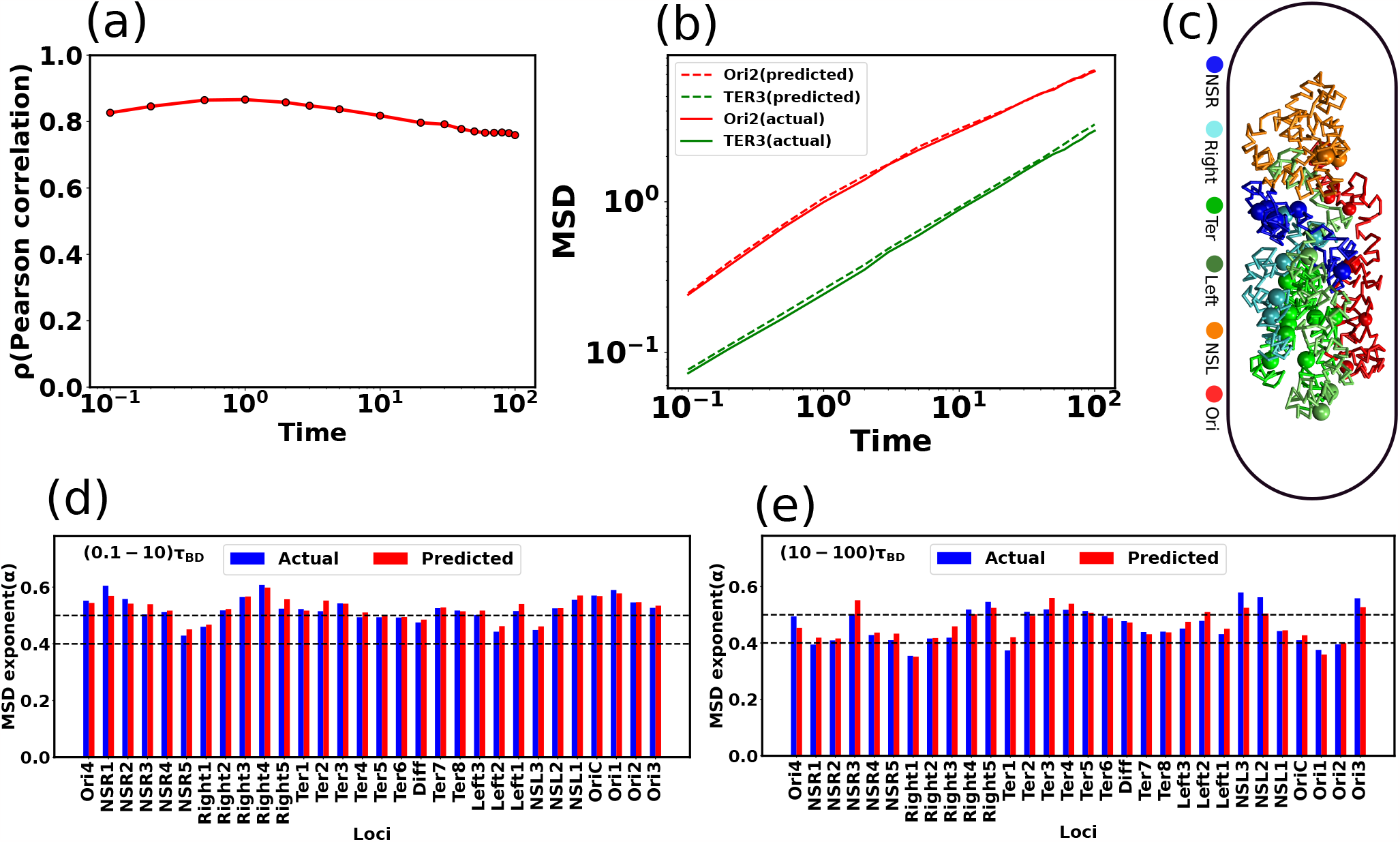
Comparison between the Actual and predicted MSD values and their exponents. (a) Pearson Correlation Coefficient (PCC) between actual and predicted MSDs a function of time. During shorter time intervals, PCC demonstrates higher values (*>* 0.8). Conversely, there is a marginal decrease in the PCC for longer time intervals. (b) Comparison of actual and predicted MSDs as a function of time for two particular loci Ori2 and Ter3. The dotted line represents the predicted MSDs, while the solid line depicts the actual MSDs. These plots illustrate a notable concordance between the observed and predicted MSDs. (c) Equilibrated snapshots of the simulated chromosome. The macrodomains are highlighted by distinct color-coded chunks, and the loci associated with each macrodomain are depicted through a spherical bead representation. Comparing the MSD exponents between observed and predicted values is illustrated for two distinct time intervals: (d) (0.1 *−*10)*τ*_*BD*_ (short time) and (e) (10*−*100)*τ*_*BD*_ (long time). Notably, all loci exhibit heterogeneous subdiffusive motion, irrespective of the time intervals. During the short time, the actual MSD exponents for all loci closely align with the predicted exponents. However, for the long time, a slight deviation in exponents between the actual and predicted values becomes apparent. (c) The PCC between the actual and predicted MSDs as a function of time for both RF and LR models. Interestingly, the LR model has much lower correlation compared to the RF model across all time scales. In the context of predicting dynamics, the RF regression model outperforms the simpler LR model. In all the plots, both the MSD and time are expressed in terms of *σ*^2^ and *τ*_*BD*_ respectively.

Now, we aim to characterize the type of diffusion for individual loci. In Figure 4(c), we present an equilibrated snapshot of the bacterial chromosome, with each color-coded chunk depicting a distinct macrodomain(MD). Each MD is comprised of various loci represented as spherical beads. We fitted the MSD values (both actual and predicted) of each individual loci with a power law:

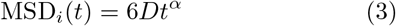

where *t, D*, and *α* are the time, diffusion constant, and exponent, respectively. Depending on the exponent *α*, one can categorize the type of diffusion; for example, *α* = 1 corresponds to normal diffusion, *α <* 1 signifies subdiffusion, and *α >* 1, indicates superdiffusion. Figures 4(d) and (e) depict the comparison of MSD exponents between actual and ML-predicted values for two different time intervals, namely (0.1 *−*10)*τ*_*BD*_ (short time) and (10 *−*100)*τ*_*BD*_ (long time), respectively. From these figures, it is evident that for the short time, the actual MSD exponents for all loci closely match the predicted exponents. However, for the long time, there is a slight deviation in exponents between the actual and predicted values. As the PCC between the actual and predicted MSD values deviates for longer times (as seen in Figure 4(a)), this discrepancy is also reflected in the exponent values. However, for both timescales, the observed and predicted dynamics exhibit subdiffusion, showcasing significant variability along the genomic coordinates, thereby indicating a heterogeneous nature of the dynamics. We found the ML-model to be robust against multiple hyperparameter (see figure S1 a)-b) and related supplemental results SR1). More importantly, the proposed ML protocol provided significantly superior dynamical prediction than a non-tree based algorithm, specifically Linear Regression (LR) (see figure S1 c) and the supplemental results SR1)

#### Identifying important chromosomal contact features crucial for loci dynamics

In general, the RF enlightens us about the quantitative extent of significance of each feature by evaluating its impact on impurity. In classification tasks, impurity is assessed through Gini impurity or information gain, while in regression, it involves variance reduction [35, 36]. During the training, within the decision trees, the greater the reduction in impurity caused by a feature, the more pivotal that feature becomes. In our RF regression model, we have a total of 928 inter-gene distance-based features. Each feature represents the distance between one particular DNA beads with all other beads. To explore the time-dependent importance of these features, we computed cumulative sums of feature importance. Figures 5(a), (b), (c), and (d) depict the cumulative sums of feature importance as a function of the total number of features for different time points: 0.1*τ*_*BD*_, 1.0*τ*_*BD*_, 10.0*τ*_*BD*_, and 100.0*τ*_*BD*_, respectively. In each plot, the vertical black dotted line highlights the number of features that contribute to 85% of the total feature importance score. We named the particular number of features as *top features*. A closer examination of the black dotted lines reveals that the number of *top features* is dynamic i.e. varying with time.

**FIG. 5.**
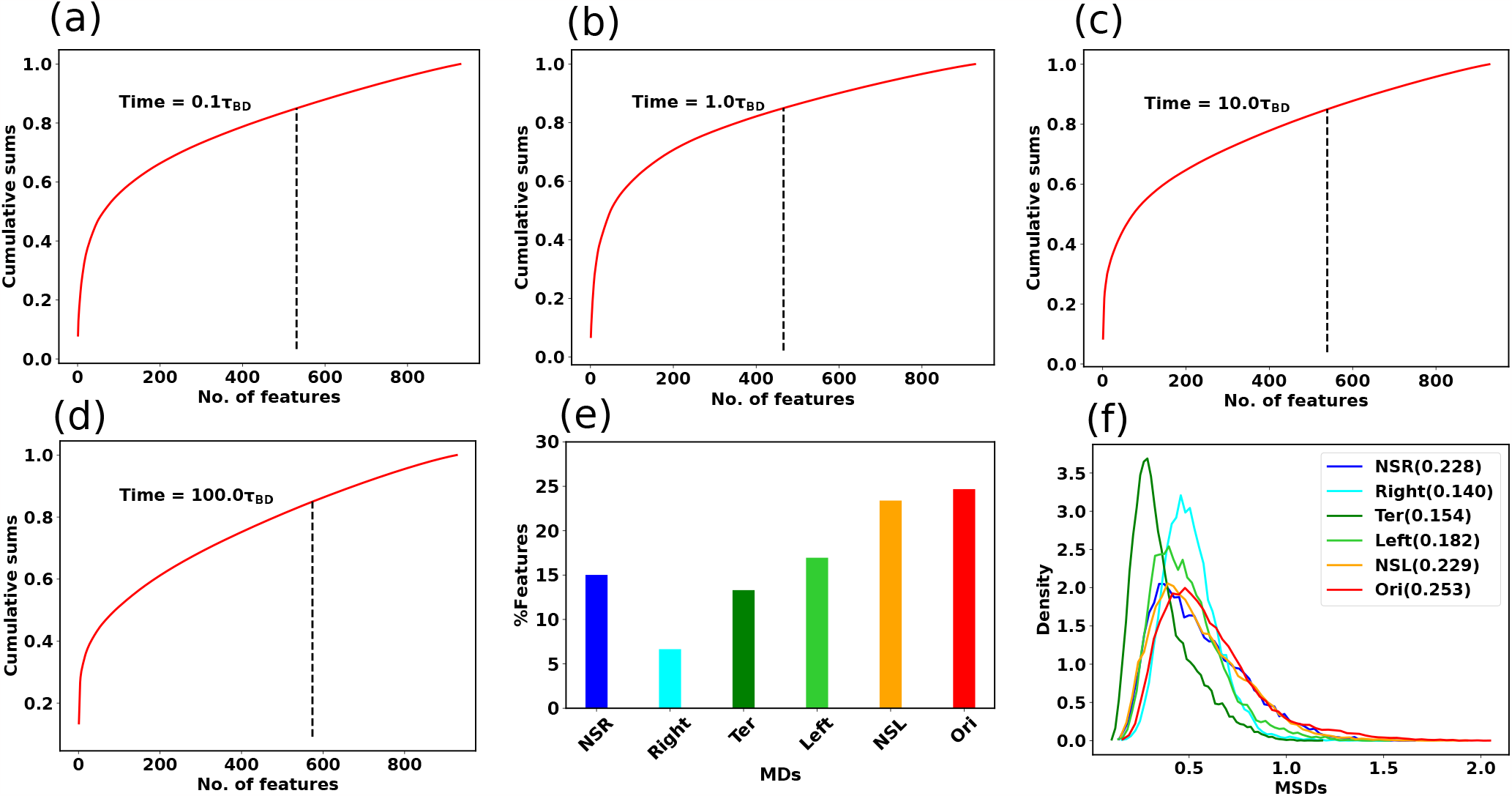
Important features specific to each macrodomain. Cumulative sums of feature importance as a function of the total number of features for different time points (a) 0.1*τ*_*BD*_, (b) 1.0*τ*_*BD*_, (c) 10.0*τ*_*BD*_, and (d) 100.0*τ*_*BD*_ respectively. Each plot includes a vertical black dotted line indicating the number of features responsible for 85% of the overall feature importance score (referred to as *top features*). The number of *top features* is varying with time. (e) The bar plot of the percentage-wise contributions of *top features* with respect to different macrodomains. Notably, Ori MD exhibits a predominant share of *top features*, while Right MD showcases a comparatively smaller proportion of these *top features*. (f) Distribution of the MSD values for all the MDs at the specific time point. The standard deviation of each distribution is reported in the legend of the plot. Notably, Ori MD shows a wider distribution compared to Right MD.

To get deeper insights into the feature importance, we have selected the *top features* at a particular time 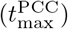 when the PCC between the actual and predicted MSDs becomes maximum. In this context, we pinpointed a total of 466 *top features* representing the 85% of the total feature important score. Within the set of 466 *top features*, we computed the percentage-wise contributions from each macrodomain. Figure 5(e) represents the bar plot of the % of *top features* with respect to different macrodomains. Quite interestingly the percentage-wise contribution of *top features* is not uniform with respect to the various macrodomains. Specifically, Ori MD exhibits the most substantial contribution, whereas Right MD demonstrates a comparatively smaller contribution. In the same spirit, we also identified the *top features* that remain common across different times, totaling 207 in number. Subsequently, we have also plotted the percentage-wise contributions of common *top features* from each macrodomain (Figure S2). The plot shows a very similar trend as Figure 5(e).

To understand this nonuniform contributions of MDs in feature importance, we have plotted the distribution of the MSD values for different MDs at the specific time point 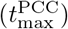. Figure 5(f) shows the distribution of the MSD values for all the MDs and the standard deviation of each distribution is reported in the legend of the plot. Notably, the plot reveals a significantly broader distribution of MSD values for Ori MD in comparison to Right MD. In the context of our machine learning model, where MSD values serve as labels for supervised learning, these findings imply that the RF regression model requires a greater number of features to construct accurate decision trees when faced with a broad distribution of training data, and conversely, fewer features are needed in the case of a narrower distribution

#### Can chromosome dynamics be reconstructed using only ML-derived important features?

For a more comprehensive grasp of functional implication of the ML-derived *top features*, we decided to consider only these particular distance-based chromosomal contact features from Hi-C map and incorporate them in our particle-based DNA model. Precisely, originally totaling 17302 Hi-C contacts, we have now reduced it to 12, 233, resulting in a notable reduction of approximately 29%.

By incorporating these subset of Hi-C contacts, we conducted a new set of simulations and compared the outcomes with our initial modelling results. We named the previous set as “actual” and the current one as “UTF” (using top features). Figure 6(a) showcases a heat map of Hi-C contact probability, where the upper and lower triangular matrices represent simulated (UTF) and experimental contact probabilities, respectively. The high PCC of 0.90 between these matrices signifies robust agreement. In terms of dynamics, we have also calculated the MSD exponent of each loci. Figures 6(b) and (c) present bar plots of the MSD exponent for all loci at two timescales: (0.1 *−*10)*τ*_*BD*_ and (10 *−*100)*τ*_*BD*_, respectively. At a shorter time scale, the MSD exponent between the actual and UTF aligns well. However, deviations emerge at longer timescales. These deviations are believed to originate from the dynamic nature of *top features*. From Figures 5(a), (b), (c), and (d), it is clear that the number of top features varies largely over time. But in our new set of simulations (UTF), we have only incorporated the *top features* related Hi-C contacts, at a particular time when the PCC between the actual and predicted MSDs becomes maximum. This modelling approach may overlook crucial features relevant for later times, impacting the accuracy of the exponent. Additionally, Figure 4(a) shows that the PCC between the actual and predicted MSDs is lower at a longer time. These effects will always provide the deviation of the exponent at a larger time scale irrespective of the time-dependent features engineering.

**FIG. 6.**
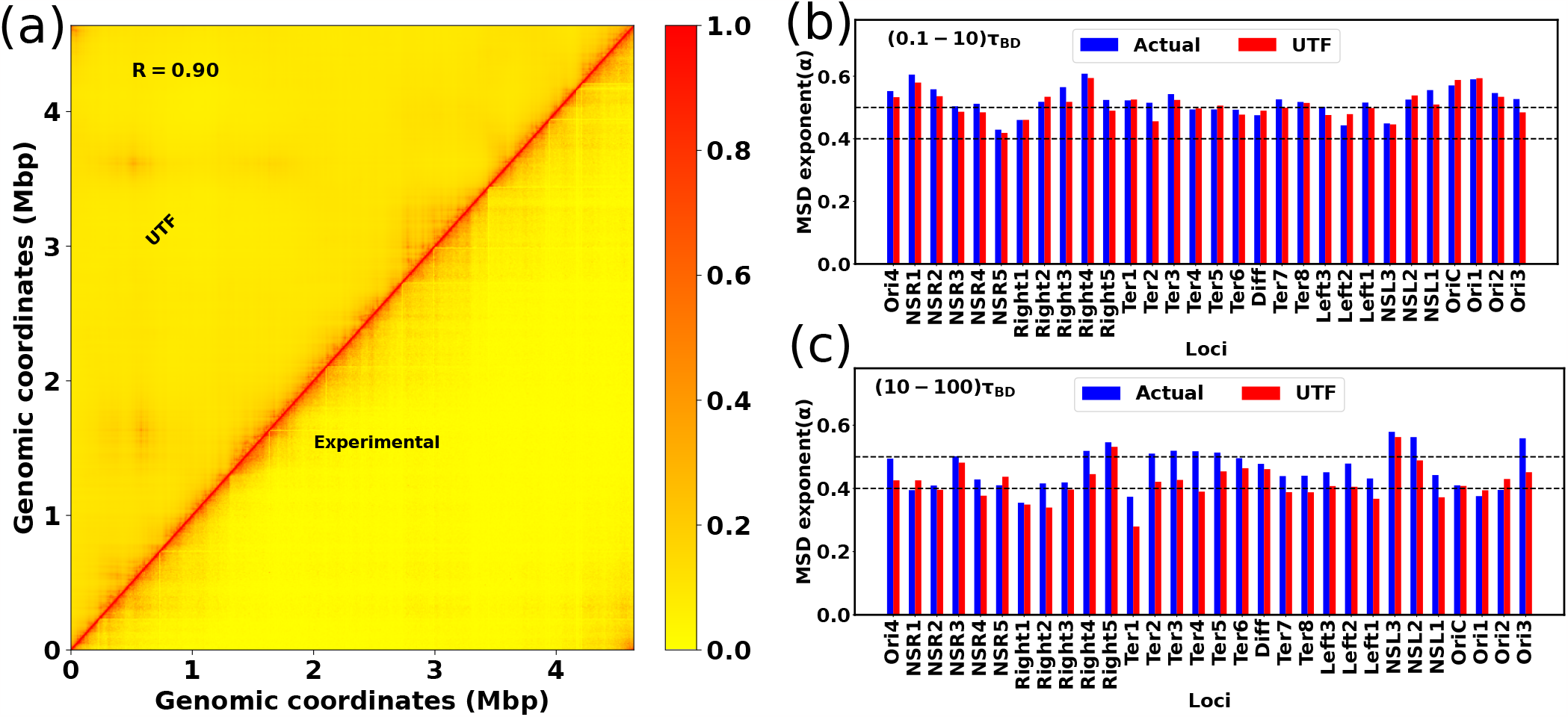
Reproduce the structure and dynamics by using important features. (a) The heat map between the experimental and simulated (UTF) contact probability matrix for WT30MM. A PCC value of 0.90 between these matrices indicates strong agreement. The bar plot of the MSD exponent for different loci at two timescales: (b) (0.1 *−*10)*τ*_*BD*_ and (c) (10 − 100)*τ*_*BD*_, respectively. At a shorter time scale, the MSD exponent between the actual and UTF matches quite well. However, at a larger time scale, they start to deviate.

Nevertheless, the deviations of the exponent are not so huge. Based on this observation, we can assert that RF regression is a powerful technique for predicting dynamics and engaging in feature engineering. The concept of important features allows us to extract the effective Hi-C contacts that can qualitatively provide the structure and dynamics.

### C. Probing prediction ability of ML-model on NAP-devoid Mutant

#### To what extent can ML recreate Hi-C matrix of Mutant chromosome?

*E. coli* intricately maintains a chromosome architecture characterized by distinct macrodomains. Several proteins are responsible for this structural management. These proteins, known as nucleoid-associated proteins (NAPs), contribute to the orchestration of chromosomal organization [39, 40]. Within this category, certain NAPs exhibit localized binding to chromosomes, while others engage in nonspecific binding. These multifaceted NAPs play discernible roles in shaping the overall organization of the chromosome. Among the NAPs, MatP stands out as a key player responsible for isolating the Ter MD from the rest of the chromosome. Specifically, MatP exhibits specific binding to 23 sites within the Ter MD, known as matS sites [11, 41]. Notably, in the absence of MatP, there is an enrichment in long-range contacts within the Ter MD and its adjacent domains [16]. Another essential protein in the realm of chromosomal structure maintenance is MukBEF, which actively facilitates long-range contacts outside the Ter MD [42]. Interestingly, when MukBEF is absent, a reduction in long-range contacts is observed across all MDs except for the Ter MD [16].

We decided to investigate the extent of feasibility of recreating Hi-C matrices for two distinct mutants, namely ΔMatP30MM and ΔMukBEF22MM, using the ML model that we had trained on wildtype (WT30MM) Hi-C map. This approach would allow us to assess the extent of intrinsic information within the WT matrix that contributes to the accuracy of reconstructing the chromosome contact map of mutants. Importantly, We do not intend to retrain the model with mutant data. Rather, the ML model aims to utilize the pre-optimized weight and bias values derived from the WT Hi-C data to generate the mutant Hi-C matrices.

In Figure 7(a) and S3(a), we compare the contact probability maps of the experimental and recreated Hi-C matrices for ΔMatP30MM and a histogram illustrating the matrix differences respectively. The substantial agreement between these matrices is emphasized by a PCC value of 0.92 and an absolute difference in mean values of 0.027. For a more detailed examination of the Hi-C matrices, we computed PCC within distinct macrodomains (MDs) (Figure 7(b)). The correlation is notably high between individual MDs and their adjacent MDs, contrasting with the lower correlation observed for MDs that are farther apart. While the experimental and recreated matrices may seem similar at first glance, a closer examination through a heat map of their differences reveals specific dissimilarities(Figure 7(c)). Notably, there is a butterfly-shaped region in the Ter and its adjacent domains, highlighted within the magenta box. These findings imply that the recreated Hi-C fails to accurately capture the interactions within the Ter and its flanking domains. Similarly, in Figure 7(d), S3(b), and (e), we depict the contact probability map, distribution of the difference in the contact matrix, and MDs-wise PCC between the experimental and recreated Hi-C matrices of ΔMukBEF22MM. The overall PCC (0.92), mean difference values(0.03), and individual PCC of each MD indicate a robust agreement between these matrices. However, a closer examination of the difference heatmap (Figure 7(f)) reveals long-range contacts across all MDs, except for the Ter MD (highlighted by the magenta box). This observation suggests that the recreated matrix fails to appropriately capture the long-range reduction of interactions for all domains except the Ter.

**FIG. 7.**
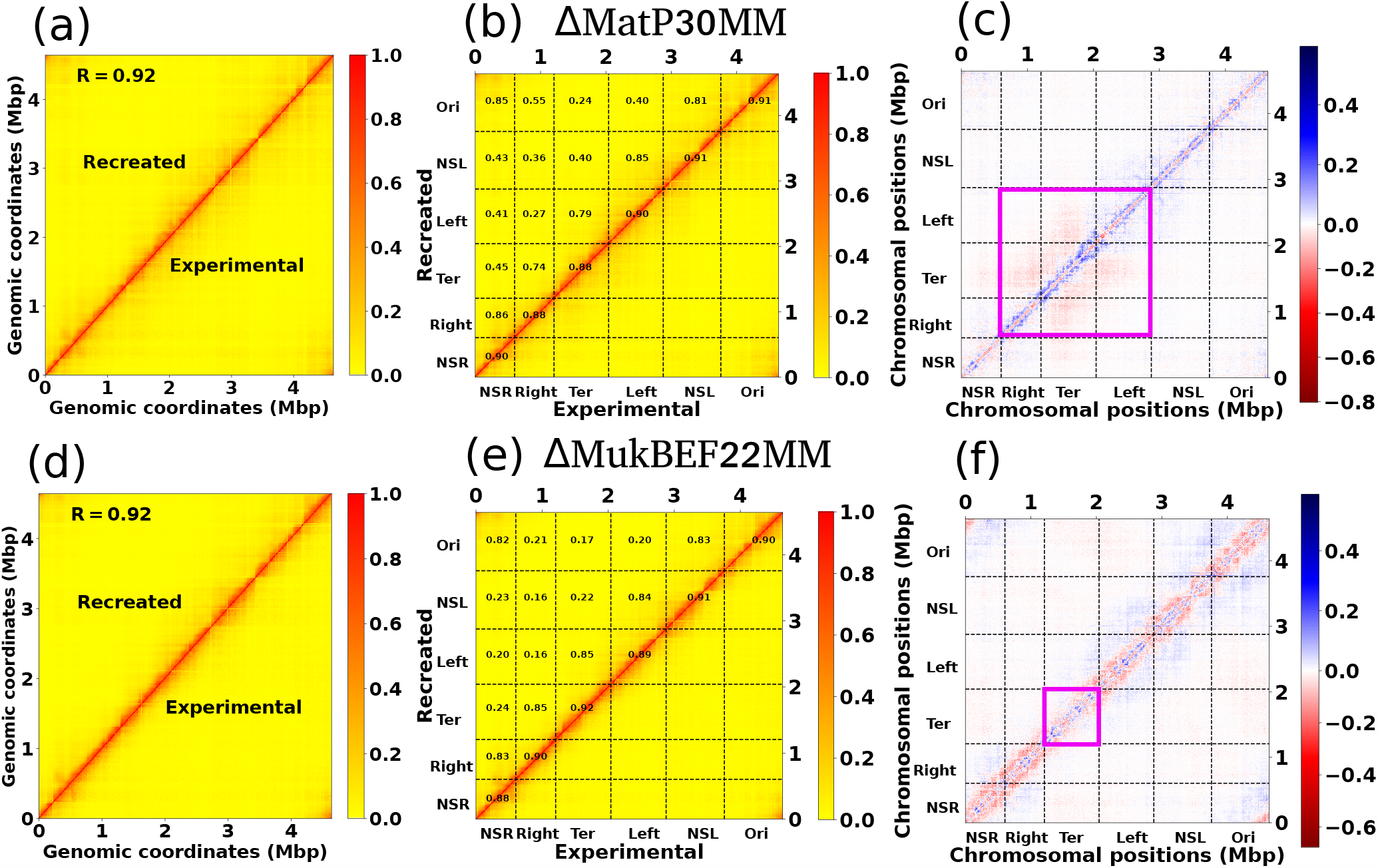
Recreation of the Hi-C matrix for different mutant by using machine learning trained model on wild type Hi-C matrix. (a) Heat map between the experimental and ML recreated contact probability matrix for ΔMatP30MM. A Pearson correlation coefficient(PCC) value of 0.92 indicates a reasonably strong agreement between them. (b) Within the heat map of the contact probability matrix, individual PCC values for each MD are presented. The PCC are notably high between individual MDs and their adjacent MDs. (c) The heat map of the difference matrix (experimental and ML recreated) reveals a butterfly-shaped region inside the Ter and its flanking domains, denoted by the magenta box. This observation suggests that the ML-recreated matrix fails to accurately capture the interactions within the Ter and its nearby domains. (d), (e), and (f) depict similar plots as in (a), (b), and (c) respectively. The only difference is that here we compare the experimental and ML recreated Hi-C matrix for ΔMukBEF22MM. From Figure (f), it is evident that there are long-range contacts across all MDs except the Ter, suggesting that the ML-derived matrix cannot accurately capture the long-range reduction of contacts for all MDs except the Ter.

The unsupervised ML model operates within the constraints of the information it has been exposed to during training, unable to generate entirely novel insights beyond its training data. Essentially, it excels at recognizing patterns within the provided datasets. Consequently, when the model is trained solely on the WT Hi-C matrix without exposure to mutant data, it inevitably falls short in accurately capturing the modified interactions specific to mutants compared to the WT type. For instance, in the recreation of the

ΔMatP30MM Hi-C matrix, the model fails to adequately capture the nuanced interactions within the Ter and its flanking MDs. Similarly, for the ΔMukBEF22MM Hi-C matrix, the model fails to accurately represent the reduction in long-range interactions for all MDs, except the Ter. Interestingly, the WT training model manages to capture various other intrinsic information crucial for recreating mutant Hi-C matrices. To support this, we have trained the same model with a random matrix whose diagonal elements are 1 and all other elements are between (0 *−*1)(*see Method section*). Attempting to reconstruct the ΔMatP30MM with this trained model resulted in an extremely poor PCC of 0.03 (Figure S4). Together these results suggest that our unsupervised ML model provides a powerful way to reconstruct the Hi-C of the mutant which is semi-quantitatively accurate.

#### How close does the ML-derived model replicate the dynamics of different mutants?

In our earlier discussion on the intrinsic structural patterns within the Hi-C matrix, we observed that the WT training model effectively captures the mutant Hi-C matrices in a semi-quantitative manner. Now, our focus is on predicting the dynamics of chromosomal loci for various mutants using the training model on WT data. In this scenario, the model utilizes previously trained “decision trees” to predict the dynamics. Figure S5 shows the PCC(*ρ*) between the actual and predicted MSDs over time for both WT and mutant (ΔMatP30MM and ΔMukBEF22MM) chromosomes. The figure indicates that the correlations for both mutants are lower compared to the WT chromosome. Additionally, the correlation for ΔMukBEF22MM deviates more from WT compared to ΔMatP30MM, suggesting that the WT training model is less accurate in capturing loci dynamics for ΔMukBEF22MM. A closer examination of the Hi-C matrices for WT and mutant cases reveals distinct patterns. For ΔMatP30MM, the contact probability in the Ter and its flanking domains deviates from the WT matrix. In contrast, for ΔMukBEF22MM, the contact probability for all macrodomains deviates from the WT matrix, except for Ter MD. This suggests that when training the model with the WT matrix, it captures information more useful for predicting the dynamics of ΔMatP30MM compared to ΔMukBEF22MM. Consequently, when predicting the chromosome dynamics for ΔMatP30MM using WT training data, the deviation in correlation is less compared to ΔMukBEF22MM.

## III. DISCUSSIONS AND SUMMARY

The organization and dynamics of the bacterial DNA are very complex and not yet fully explored. To address this, the utilization of the Hi-C integrated [21–24, 43] and cross-linked [44, 45] based polymer model offers a versatile means of exploring the organization and dynamics of *E. coli* DNA. In this study, we present a comprehensive approach, employing a set of machine learning (ML) algorithms to gain insights into the structural and dynamical aspects of bacterial chromosome. By leveraging a combination of Autoencoder-based structural analysis, and Random Forest(RF) regression for predicting chromosomal dynamics, our work provides valuable insights into an intricate pattern of bacterial chromosomal organization and it emergent dynamics.

In the first part of the study, we mainly focused on extracting essential structural information, that is hidden under-neath the Hi-C matrix, using an unsupervised deep neural network known as Autoencoder. The low-dimensional representation of the Hi-C data interestingly identifies chromosomal macrodomains (MDs) as key structural pattern in an automated way (Figures 2). Notably, the comparison between MDs derived from our ML model and those experimentally identified reveals a high correlation, suggesting meaningful connections between the ML-derived insights and real-world biological phenomena. Moreover, when recreating the Hi-C matrix for various mutants using wild-type (WT) training data (Figures 7), our model demonstrates the ability to capture diverse intrinsic information crucial for reconstructing mutant Hi-C matrices. Significantly, the observed structural properties closely align with established experimental findings, underscoring the effectiveness of our approach in capturing biologically relevant phenomena.

After delving into the structural insights extracted from the Hi-C data, the second part of our study focused on harnessing the power of ML to predict the crucial subsets of chromosomal contact features that can optimally explain the previously reported heterogeneous subdiffusion of chromosomal loci. We employed a Hi-C embedded polymer model for the *E. coli* chromosome, representing short and long-range Hi-C contacts as effective springs with spring constants dependent on contact probabilities. Subsequently, we utilized Random Forest(RF) regression, a powerful supervised machine learning algorithm, to predict the dynamics and MSD exponent of individual chromosomal loci. Our results demonstrated a high correlation between the predicted and actual MSD values for shorter time scales (Figures 4). These observations highlight the efficacy of our machine learning model in capturing the complex relationship between structural features and dynamic behavior. However, at a large time scale, the correlation between the actual and predicted values decreases. We hypothesized that at larger time scales, the system loses its initial structural information, leading to a reduction in correlation. Despite this decrease, it remained relatively modest. Additionally, the comparison between Random Forest Regression and Linear Regression (LR) underscored the superiority of the former in our predictive task. In a simple LR method, the model aims to establish an approximate linear relationship between the dependent variable (the target) and independent variables (features) by minimizing the sum of squared residuals. The poor performance of the linear regression model suggested that pairwise distance is not a trivial feature for predicting dynamics. These observations imply a lack of a direct one-to-one correspondence between pairwise distance and dynamics; rather, complex relationships exist between them, successfully captured by RF regression for a more accurate prediction of dynamics.

We systematically evaluated the robustness of our RF regression model by varying hyperparameters (Figures S1). The consistent performance across different hyperparameter values affirmed the reliability of our machine-learning approach. An essential aspect of our study involved extracting *top features* through feature importance analysis, providing valuable insights into the critical elements influencing chromosomal dynamics. However, these *top features* are dynamic rather than static, exhibiting variations over time (Figures 5(a)-(d)). The distribution of *top features* at a particular time across different macrodomains revealed non-uniform contributions (Figure 5(e)). Particularly, the Ori macrodomain exhibited a more substantial contribution compared to the Right macrodomain. This non-uniformity was further elucidated by examining the distribution of MSD values for different macrodomains, with Ori displaying a broader distribution (Figure 5(f)). These findings indicate that the model requires more features to accurately capture dynamics when faced with a broader distribution of training data. Moreover, by incorporating the *top features* associated with Hi-C contacts into a polymer-based model, we can effectively reconstruct both the experimental Hi-C matrix and the dynamical behavior of chromosomal loci at a short time scale (Figures 6). These findings strongly imply the utility of our RF regression model for feature engineering, specifically in extracting the important Hi-C contacts that play a crucial role in influencing chromosomal dynamics. However, our model does have limitations; for instance, it may not fully capture interactions specific to mutants as it is trained solely on wild-type data.

Our work enhances the broader understanding of bacterial chromosome via computational modeling with ML techniques. The identification of macrodomains and the prediction of chromosomal dynamics offer a comprehensive view of the intricate interplay between structure and function in bacterial genomes. However, our approach can be extended beyond the realm of bacterial chromosomes. We are optimistic about the broader applicability of our methodologies in addressing more complex systems, including proteins and glass, to extract structural insights and predict dynamics. The eukaryotic cells, which possess multiple chromosomes, have been subjects of ML-driven studies [27–30] such as: subcompartment annotation of the genome [27, 28] by using inter-chromosomal contact, enhancing the resolution of Hi-C data [46, 47]. In the realm of glassy systems, there has recently been a plethora of studies predicting dynamics using a combination of structural properties and diverse machine-learning algorithms [48– 55]. While many of these studies leverage a multitude of structural features for dynamic predictions, we hope that our approach may offer an effective avenue for predicting dynamics and facilitating feature engineering.

## METHODS

### The training of the Autoencoder

The Autoen-coder in our study consists of nine fully connected sequential layers. We have used a single Hi-C contact probability matrix for training the Autoencoder. We utilize the Leaky rectied linear unit (ReLU) activation function for each layer, except for the last layer. Given that our Hi-C contacts fall within the range of 0 to 1, the sigmoid activation function is applied to the final layer. To optimize the weights and biases of each node, we utilize the Adam optimizer [56] to minimize the loss function. We choose binary cross-entropy (BCE) [27, 57] as a loss function which is defined as

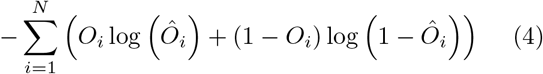

where *Ô*, and *O* are the model output, target output, and *N* = 928 respectively. The Autoencoder is trained with a batch size of 30 for input data and a learning rate of 0.001. Notably, in training the Autoencoder on the Random matrix, we have used the same architecture, modifying only the dimension of the latent space *L*_*d*_. Our observations indicate that achieving a Pearson Correlation Coefficient (PCC) value of 0.79 between the actual random matrix and the Autoencoder-derived matrix necessitates setting *L*_*d*_ to 40 (Figure S6). All aspects related to the Autoencoder, including training and implementation, are conducted using the Python implementation of Tensorflow [58] and Keras [59].

### Simulation model details

We have applied our previously established bead-spring model [21, 22] for integrating Hi-C data in *E. coli* chromosome, where each bead corresponds to 5 *×* 10^3^ bp (5 kb). The harmonic interaction between adjacent beads is governed by a spring constant of *k*_*spring*_ = 300*k*_*B*_*T/σ*^2^, where *σ* denotes the bead diameter. The inclusion of Hi-C contacts in the polymer chain introduces an effective spring with a spring constant dependent on contact probabilities. This process involves: (i) transforming the Hi-C probability matrix into a distance matrix using the formula:

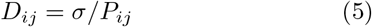

where *i* and *j* are the row and column index of the matrix respectively. (ii) By using *D*_*ij*_, we have calculated the effective spring constant as:

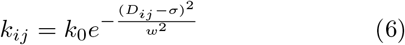

Here, *k*_0_ serves as the upper bound of the spring constant, and *w*^2^ is a constant value. In our simulation, we have maintained the values of *k*_0_ and *w*^2^ consistent with our previous study [21, 22], specifically *k*_0_ = 10*k*_*B*_*T/σ*^2^ and *w*^2^ = 0.3. The potential related to Hi-C contacts, denoted as *E*_*Hi−C*_(*r*_*ij*_), is expressed as:

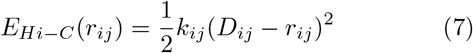

In our simulation, contacts with a spring constant *k*_*ij*_ *<* 10^*−*7^ are disregarded to avoid unnecessary low-value contacts. additionally, the nonbonded interactions between beads are modeled using the repulsive part of the Lenard-Jones potential, i.e., 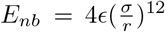. All particles are confined within a spherocylinder, mimicking the cell wall, with a length of *L* = 45.754*σ* and diameter *d* = 12.181*σ*. The confinement potential is defined as:

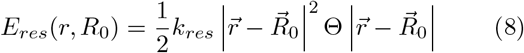

Here, *R*_0_ represents the center of the spherocylinder, and *k*_*res*_ is the spring constant controlling confinement softness (set to 310*k*_*B*_*T/σ*^2^). The step function Θ activates if any particle surpasses the confinement boundaries. The total Hamiltonian of the system is given by:

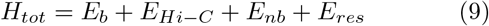

Here, *E*_*b*_, *E*_*Hi−C*_, *E*_*nb*_, and *E*_*res*_ represent the potentials for bonded, Hi-C restraining, non-bonded, and confinement restraining interactions, respectively. All the simulations were conducted using an modified version of open-source software GROMACS 5.0.6 [60], while for the implementation of random forest regression, we utilized the Python library known as scikit-learn [61].

## CODE AVAILABILITY

The code and accompanying documentation for training the Autoencoder and Random Forest regression model can be accessed through GitHub at the following URL: https://github.com/palash892/Hi-C_ML_structure-dynamics

## ACKNOWLEDGMENTS

All the authors acknowledge Tata Institute of Fundamental Research Hyderabad, India for providing the support of computing resources. We acknowledge support of the Department of Atomic Energy, Government of India, under Project Identification No. RTI 4007

## References

[1] Benjamin Volkmer and Matthias Heinemann. Condition-dependent cell volume and concentration of escherichia coli to facilitate data conversion for systems biology modeling. PloS one, 6(7):e23126, 2011.

[2] G Reshes, S Vanounou, I Fishov, and M Feingold. Timing the start of division in e. coli: a single-cell study. Physical biology, 5(4):046001, 2008.

[3] David C Grainger, Douglas Hurd, Marcus Harrison, Jolyon Holdstock, and Stephen JW Busby. Studies of the distribution of escherichia coli camp-receptor protein and rna polymerase along the e. coli chromosome. Proceedings of the National Academy of Sciences, 102(49):17693–17698, 2005.

[4] Paul A Wiggins, Keith C Cheveralls, Joshua S Martin, Robert Lintner, and Jané Kondev. Strong intranucleoid interactions organize the escherichia coli chromosome into a nucleoid filament. Proceedings of the National Academy of Sciences, 107(11):4991–4995, 2010.

[5] Anjana Badrinarayanan, Rodrigo Reyes-Lamothe, Stephan Uphoff, Mark C Leake, and David J Sherratt. In vivo architecture and action of bacterial structural maintenance of chromosome proteins. Science, 338(6106):528–531, 2012.

[6] Somenath Bakshi, Albert Siryaporn, Mark Goulian, and James C Weisshaar. Superresolution imaging of ribosomes and rna polymerase in live escherichia coli cells. Molecular microbiology, 85(1):21–38, 2012.

[7] Hironori Niki, Yoshiharu Yamaichi, and Sota Hiraga. Dynamic organization of chromosomal dna in escherichia coli. Genes & Development, 14(2):212–223, 2000.

[8] Michèle Valens, Stéphanie Penaud, Michèle Rossignol, François Cornet, and Frédéric Boccard. Macrodomain organization of the escherichia coli chromosome. The EMBO journal, 23(21):4330–4341, 2004.

[9] Olivier Espéli and Frédéric Boccard. Organization of the escherichia coli chromosome into macrodomains and its possible functional implications. Journal of structural biology, 156(2):304–310, 2006.

[10] Olivier Espeli, Romain Mercier, and Frédéric Boccard. Dna dynamics vary according to macrodomain topography in the e. coli chromosome. Molecular microbiology, 68(6):1418–1427, 2008.

[11] Romain Mercier, Marie-Agnès Petit, Sophie Schbath, Stephane Robin, Meriem El Karoui, Frédéric Boccard, and Olivier Espéli. The matp/mats site-specific system organizes the terminus region of the e. coli chromosome into a macrodomain. Cell, 135(3):475–485, 2008.

[12] Sonja Julia Messerschmidt and Torsten Waldminghaus. Dynamic organization: chromosome domains in escherichia coli. Journal of molecular microbiology and biotechnology, 24(5-6):301–315, 2015.

[13] Job Dekker, Karsten Rippe, Martijn Dekker, and Nancy Kleckner. Capturing chromosome conformation. science, 295(5558):1306–1311, 2002.

[14] Josée Dostie, Todd A Richmond, Ramy A Arnaout, Rebecca R Selzer, William L Lee, Tracey A Honan, Eric D Rubio, Anton Krumm, Justin Lamb, Chad Nusbaum, et al. Chromosome conformation capture carbon copy (5c): a massively parallel solution for mapping interactions between genomic elements. Genome research, 16(10):1299–1309, 2006.

[15] Erez Lieberman-Aiden, Nynke L Van Berkum, Louise Williams, Maxim Imakaev, Tobias Ragoczy, Agnes Telling, Ido Amit, Bryan R Lajoie, Peter J Sabo, Michael O Dorschner, et al. Comprehensive mapping of long-range interactions reveals folding principles of the human genome. science, 326(5950):289–293, 2009.

[16] Virginia S Lioy, Axel Cournac, Martial Marbouty, Stéphane Duigou, Julien Mozziconacci, Olivier Espéli, Frédéric Boccard, and Romain Koszul. Multiscale structuring of the e. coli chromosome by nucleoid-associated and condensin proteins. Cell, 172(4):771–783, 2018.

[17] Stephanie C Weber, Andrew J Spakowitz, and Julie A Theriot. Bacterial chromosomal loci move subdiffusively through a viscoelastic cytoplasm. Physical review letters, 104(23):238102, 2010.

[18] Stephanie C Weber, Julie A Theriot, and Andrew J Spakowitz. Subdiffusive motion of a polymer composed of subdiffusive monomers. Physical Review E, 82(1):011913, 2010.

[19] Avelino Javer, Zhicheng Long, Eileen Nugent, Marco Grisi, Kamin Siriwatwetchakul, Kevin D Dorfman, Pietro Cicuta, and Marco Cosentino Lagomarsino. Short-time movement of e. coli chromosomal loci depends on coordinate and subcellular localization. Nature communications, 4(1):3003, 2013.

[20] Stephanie C Weber, Andrew J Spakowitz, and Julie A Theriot. Nonthermal atp-dependent fluctuations contribute to the in vivo motion of chromosomal loci. Proceedings of the National Academy of Sciences, 109(19):7338–7343, 2012.

[21] Abdul Wasim, Ankit Gupta, and Jagannath Mondal. A hi–c data-integrated model elucidates e. coli chromosomes multiscale organization at various replication stages. Nucleic acids research, 49(6):3077–3091, 2021.

[22] Palash Bera, Abdul Wasim, and Jagannath Mondal. Hi-c embedded polymer model of escherichia coli reveals the origin of heterogeneous subdiffusion in chromosomal loci. Physical Review E, 105(6):064402, 2022.

[23] Abdul Wasim, Ankit Gupta, Palash Bera, and Jagannath Mondal. Interpretation of organizational role of proteins on e. coli nucleoid via hi-c integrated model. Biophysical Journal, 122(1):63–81, 2023.

[24] Abdul Wasim, Palash Bera, and Jagannath Mondal. Development of a data-driven integrative model of a bacterial chromosome. Journal of Chemical Theory and Computation, 0(0):null, 2023. PMID: 37083406.

[25] Bogdan Bintu, Leslie J Mateo, Jun-Han Su, Nicholas A Sinnott-Armstrong, Mirae Parker, Seon Kinrot, Kei Yamaya, Alistair N Boettiger, and Xiaowei Zhuang. Super-resolution chromatin tracing reveals domains and cooperative interactions in single cells. Science, 362(6413):eaau1783, 2018.

[26] Suhas SP Rao, Miriam H Huntley, Neva C Durand, Elena K Stamenova, Ivan D Bochkov, James T Robinson, Adrian L Sanborn, Ido Machol, Arina D Omer, Eric S Lander, et al. A 3d map of the human genome at kilo-base resolution reveals principles of chromatin looping. Cell, 159(7):1665–1680, 2014.

[27] Kyle Xiong and Jian Ma. Revealing hi-c subcompartments by imputing inter-chromosomal chromatin interactions. Nature communications, 10(1):5069, 2019.

[28] Haitham Ashoor, Xiaowen Chen, Wojciech Rosikiewicz, Jiahui Wang, Albert Cheng, Ping Wang, Yijun Ruan, and Sheng Li. Graph embedding and unsupervised learning predict genomic sub-compartments from hic chromatin interaction data. Nature communications, 11(1):1173, 2020.

[29] Pascal Vincent, Hugo Larochelle, Yoshua Bengio, and Pierre-Antoine Manzagol. Extracting and composing robust features with denoising autoencoders. In Proceedings of the 25th international conference on Machine learning, pages 1096–1103, 2008.

[30] Geoff Fudenberg, David R Kelley, and Katherine S Pollard. Predicting 3d genome folding from dna sequence with akita. Nature methods, 17(11):1111–1117, 2020.

[31] Cheng-Yuan Liou, Wei-Chen Cheng, Jiun-Wei Liou, and Daw-Ran Liou. Autoencoder for words. Neurocomputing, 139:84–96, 2014.

[32] Junhai Zhai, Sufang Zhang, Junfen Chen, and Qiang He. Autoencoder and its various variants. In 2018 IEEE international conference on systems, man, and cybernetics (SMC), pages 415–419. IEEE, 2018.

[33] Aristidis Likas, Nikos Vlassis, and Jakob J Verbeek. The global k-means clustering algorithm. Pattern recognition, 36(2):451–461, 2003.

[34] Trupti M Kodinariya, Prashant R Makwana, et al. Review on determining number of cluster in k-means clustering. International Journal, 1(6):90–95, 2013.

[35] Leo Breiman. Random forests. Machine learning, 45:5–32, 2001.

[36] Leo Breiman. Classification and regression trees. Routledge, 2017.

[37] Asaph Widmer-Cooper, Peter Harrowell, and H Fynewever. How reproducible are dynamic heterogeneities in a supercooled liquid? Physical review letters, 93(13):135701, 2004.

[38] Asaph Widmer-Cooper and Peter Harrowell. On the study of collective dynamics in supercooled liquids through the statistics of the isoconfigurational ensemble. The Journal of chemical physics, 126(15), 2007.

[39] Martijn S Luijsterburg, Maarten C Noom, Gijs JL Wuite, and Remus Th Dame. The architectural role of nucleoid-associated proteins in the organization of bacterial chromatin: a molecular perspective. Journal of structural biology, 156(2):262–272, 2006.

[40] Talukder Ali Azam and Akira Ishihama. Twelve species of the nucleoid-associated protein from escherichia coli: sequence recognition specificity and dna binding affinity. Journal of Biological Chemistry, 274(46):33105–33113, 1999.

[41] Pauline Dupaigne, Nam K Tonthat, Olivier Espéli, Travis Whitfill, Frédéric Boccard, and Maria A Schumacher. Molecular basis for a protein-mediated dna-bridging mechanism that functions in condensation of the e. coli chromosome. Molecular cell, 48(4):560–571, 2012.

[42] Sophie Nolivos and David Sherratt. The bacterial chromosome: architecture and action of bacterial smc and smc-like complexes. FEMS microbiology reviews, 38(3):380–392, 2014.

[43] Joris JB Messelink, Muriel CF van Teeseling, Jacqueline Janssen, Martin Thanbichler, and Chase P Broedersz. Learning the distribution of single-cell chromosome conformations in bacteria reveals emergent order across genomic scales. Nature communications, 12(1):1963, 2021.

[44] Srikanth Subramanian and Seàn M Murray. Subdiffusive movement of chromosomal loci in bacteria explained by dna bridging. Physical Review Research, 5(2):023034, 2023.

[45] Tejal Agarwal, GP Manjunath, Farhat Habib, and Apratim Chatterji. Bacterial chromosome organization. ii. few special cross-links, cell confinement, and molecular crowders play the pivotal roles. The Journal of Chemical Physics, 150(14), 2019.

[46] Yan Zhang, Lin An, Jie Xu, Bo Zhang, W Jim Zheng, Ming Hu, Jijun Tang, and Feng Yue. Enhancing hi-c data resolution with deep convolutional neural network hicplus. Nature communications, 9(1):750, 2018.

[47] Hao Hong, Shuai Jiang, Hao Li, Guifang Du, Yu Sun, Huan Tao, Cheng Quan, Chenghui Zhao, Ruijiang Li, Wanying Li, et al. Deephic: A generative adversarial network for enhancing hi-c data resolution. PLoS computational biology, 16(2):e1007287, 2020.

[48] Victor Bapst, Thomas Keck, A Grabska-Barwińska, Craig Donner, Ekin Dogus Cubuk, Samuel S Schoenholz, Annette Obika, Alexander WR Nelson, Trevor Back, Demis Hassabis, et al. Unveiling the predictive power of static structure in glassy systems. Nature Physics, 16(4):448–454, 2020.

[49] Emanuele Boattini, Susana Marín-Aguilar, Saheli Mitra, Giuseppe Foffi, Frank Smallenburg, and Laura Filion. Autonomously revealing hidden local structures in supercooled liquids. Nature communications, 11(1):5479, 2020.

[50] Emanuele Boattini, Frank Smallenburg, and Laura Filion. Averaging local structure to predict the dynamic propensity in supercooled liquids. Physical Review Letters, 127(8):088007, 2021.

[51] Rinske M Alkemade, Frank Smallenburg, and Laura Filion. Improving the prediction of glassy dynamics by pin-pointing the local cage. The Journal of Chemical Physics, 158(13), 2023.

[52] Rinske M Alkemade, Emanuele Boattini, Laura Filion, and Frank Smallenburg. Comparing machine learning techniques for predicting glassy dynamics. The Journal of Chemical Physics, 156(20), 2022.

[53] Hayato Shiba, Masatoshi Hanai, Toyotaro Suzumura, and Takashi Shimokawabe. Botan: Bond targeting network for prediction of slow glassy dynamics by machine learning relative motion. The Journal of Chemical Physics, 158(8), 2023.

[54] Samuel S Schoenholz, Ekin D Cubuk, Daniel M Sussman, Efthimios Kaxiras, and Andrea J Liu. A structural approach to relaxation in glassy liquids. Nature Physics, 12(5):469–471, 2016.

[55] Gerhard Jung, Giulio Biroli, and Ludovic Berthier. Predicting dynamic heterogeneity in glass-forming liquids by physics-inspired machine learning. Physical Review Letters, 130(23):238202, 2023.

[56] Diederik P Kingma and Jimmy Ba. Adam: A method for stochastic optimization. arXiv preprint arXiv:1412.6980, 2014.

[57] Antonia Creswell, Kai Arulkumaran, and Anil A Bharath. On denoising autoencoders trained to minimise binary cross-entropy. arXiv preprint arXiv:1708.08487, 2017.

[58] Martín Abadi, Ashish Agarwal, Paul Barham, Eugene Brevdo, Zhifeng Chen, Craig Citro, Greg S Corrado, Andy Davis, Jeffrey Dean, Matthieu Devin, et al. Tensorflow: Large-scale machine learning on heterogeneous distributed systems. arXiv preprint arXiv:1603.04467, 2016.

[59] Ekaba Bisong and Ekaba Bisong. Tensorflow 2.0 and keras. Building Machine Learning and Deep Learning Models on Google Cloud Platform: A Comprehensive Guide for Beginners, pages 347–399, 2019.

[60] Mark James Abraham, Teemu Murtola, Roland Schulz, Szilàrd Pàll, Jeremy C Smith, Berk Hess, and Erik Lindahl. Gromacs: High performance molecular simulations through multi-level parallelism from laptops to super-computers. SoftwareX, 1:19–25, 2015.

[61] Fabian Pedregosa, Gaël Varoquaux, Alexandre Gramfort, Vincent Michel, Bertrand Thirion, Olivier Grisel, Mathieu Blondel, Peter Prettenhofer, Ron Weiss, Vincent Dubourg, et al. Scikit-learn: Machine learning in python. the Journal of machine Learning research, 12:2825–2830, 2011.

