## Supplementary material for "Machine Learning Unravels Inherent Structural Patterns in *Escherichia coli* Hi-C Matrices and Predicts DNA Dynamics": Suplemental tables and figures

#### 0.1 Confusion Matrix

In Figure 1, we present the tabular representation of the confusion matrix. In this context, we have computed various classification metrics applicable to both binary and multi-class systems.

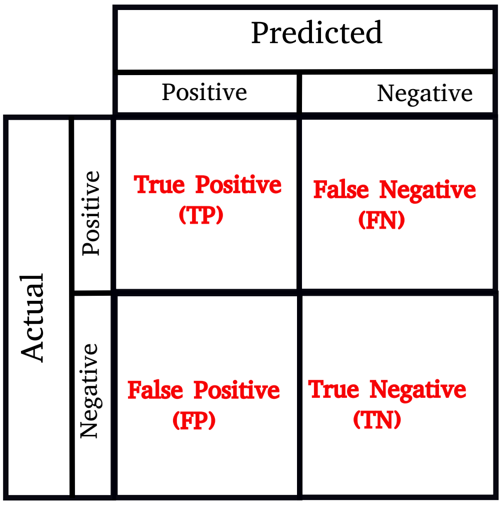


Figure 1: The architecture of the confusion matrix. The various classification metrics such as Accuracy, Precision, Recall, and F1-Score can be derived from this confusion matrix.

| | Predicted | | | | | |
|---|---|---|---|---|---|---|
| Actual | A | B | C | D | E | F |
| A | $TP_A$ | $E_{AB}$ | $E_{AC}$ | $E_{AD}$ | $E_{AE}$ | $E_{AF}$ |
| B | $E_{BA}$ | $TP_B$ | $E_{BC}$ | $E_{BD}$ | $E_{BE}$ | $E_{BF}$ |
| C | $E_{CA}$ | $E_{CB}$ | $TP_C$ | $E_{CD}$ | $E_{CE}$ | $E_{CF}$ |
| D | $E_{DA}$ | $E_{DB}$ | $E_{DC}$ | $TP_D$ | $E_{DE}$ | $E_{DF}$ |
| E | $E_{EA}$ | $E_{EB}$ | $E_{EC}$ | $E_{ED}$ | $TP_E$ | $E_{EF}$ |
| F | $E_{FA}$ | $E_{FB}$ | $E_{FC}$ | $E_{FD}$ | $E_{FE}$ | $TP_F$ |

Table 1: **Confusion matrix for six class system**

Various classification metrics are commonly defined as follows:

$$\text{Accuracy} = \frac{TP + TN}{TP + TN + FP + FN}$$

$$\text{Precision} = \frac{TP}{TP + FP}$$

$$\text{Recall} = \frac{TP}{TP + FN}$$

$$\text{F1-Score} = 2 \times \frac{\text{Precision} \times \text{Recall}}{\text{Precision} + \text{Recall}}$$

For many class system, one can define the overall Accuracy, class wise Precision, Recall, and F1-Scoreas as follows:

$$\text{Accuracy} = \frac{\text{Total correct classification}}{\text{All classification}} = \frac{TP_A + TP_B + TP_C + TP_D + TP_E + TP_F}{TP_A + TP_B + TP_C + TP_D + TP_E + TP_F + E_{AB} + ... + E_{FE}}$$

$$P_A = \frac{TP_A}{TP_A + E_{BA} + E_{CA} + E_{DA} + E_{EA} + E_{FA}}$$

$$\vdots$$

$$P_F = \frac{TP_F}{TP_F + E_{AF} + E_{BF} + E_{CF} + E_{DF} + E_{EF}}$$

$$R_A = \frac{TP_A}{TP_A + E_{AB} + E_{AC} + E_{AD} + E_{AE} + E_{AF}}$$

$$\vdots$$

$$R_F = \frac{TP_F}{TP_F + E_{FA} + E_{FB} + E_{FC} + E_{FD} + E_{FE}}$$

$$\text{F1-Score}_A = 2 \times \frac{P_A \times R_A}{P_A + R_A}$$

$$\vdots$$

$$\text{F1-Score}_F = 2 \times \frac{P_F \times R_F}{P_F + R_F}$$

Here $P_A ... P_F$, $R_A ... R_F$, F1-Score$_A$...F1-Score$_F$ represent the Precision, Recall, and F1-Score for different classes respectively.

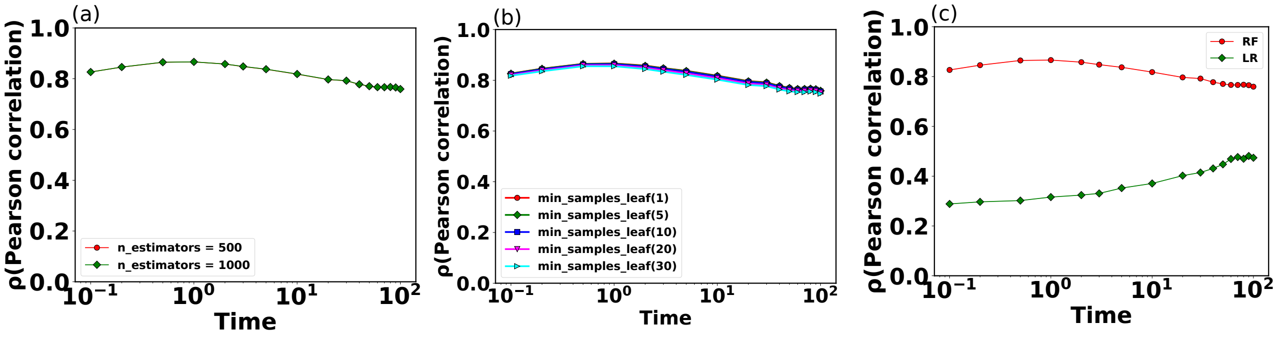


Figure S1: **Robustness of Random Forest (RF) regression and comparison with Linear Regression (LR).** The Pearson Correlation Coefficient (PCC) is depicted as a function of time for varying values of (a) n_estimators and (b) min_samples_leaf, respectively. Notably, our RF regression model has been configured with (n_estimators = 500 and min_samples_leaf = 1). The PCC exhibits minimal variation across different choices of these hyperparameters, suggesting the robustness of our model, especially within the chosen parameter set. (c) The PCC between the actual and predicted MSDs as a function of time for both RF and LR models. Interestingly, the LR model has much lower correlation compared to the RF model across all time scales. In the context of predicting dynamics, the RF regression model outperforms the simpler LR model. In all the plots, the time is expressed in terms of $\tau_{BD}$.

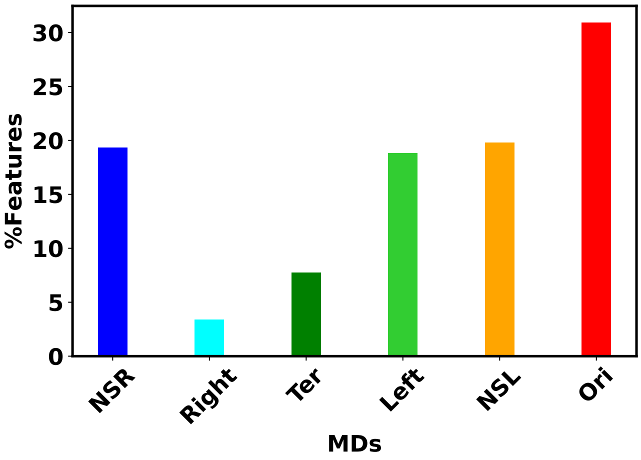


Figure S2: The bar plot of the percentage-wise contributions of common *top features* with respect to different macrodomains. Notably, Ori MD exhibits a predominant share of *top features*, while Right MD showcases a comparatively smaller proportion of these common *top features*.

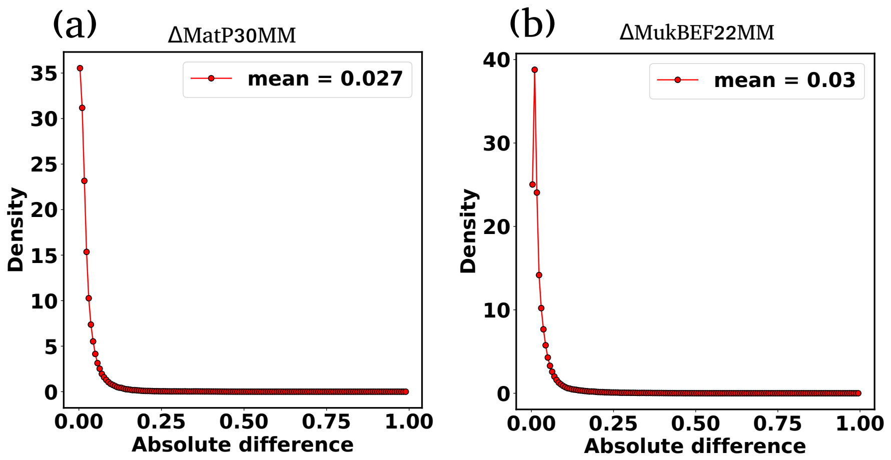


Figure S3: (a)The distribution of the absolute difference between the experimental and ML-recreated contact probability matrices for ΔMatP30MM. (b) The distribution of the absolute difference between the experimental and ML-recreated contact probability matrices for ΔMukBEF22MM.

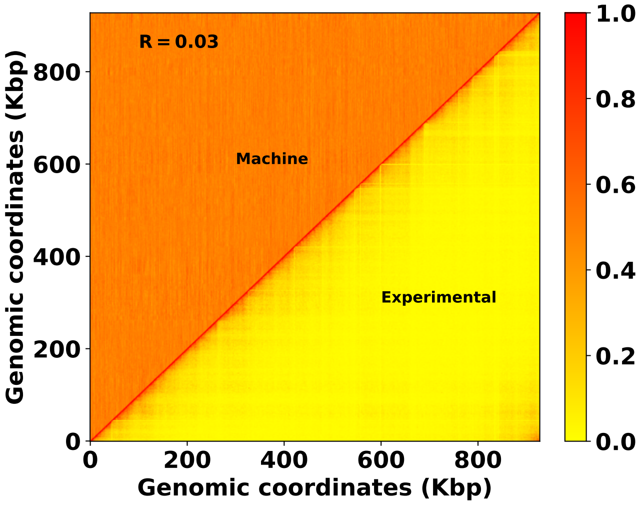


Figure S4: Comparison of experimental and ML recreated Hi-C matrix for ΔMatP30MM. We recreated the Hi-C matrix using the trained model on random matrix. The notably low value of the Pearson Correlation Coefficient (PCC) implies a poor recreation.

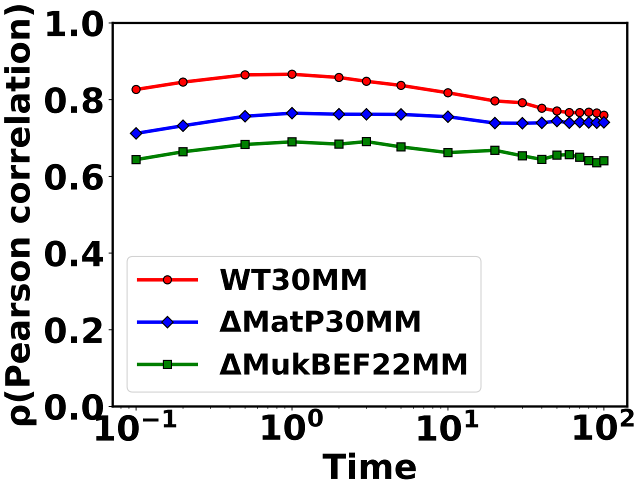


Figure S5: Pearson correlation coefficient (PCC) ($\rho$) between actual and predicted mean squared displacements (MSDs) over time for both WT and mutants ($\Delta$MatP30MM and $\Delta$MukBEF22MM) chromosomes.

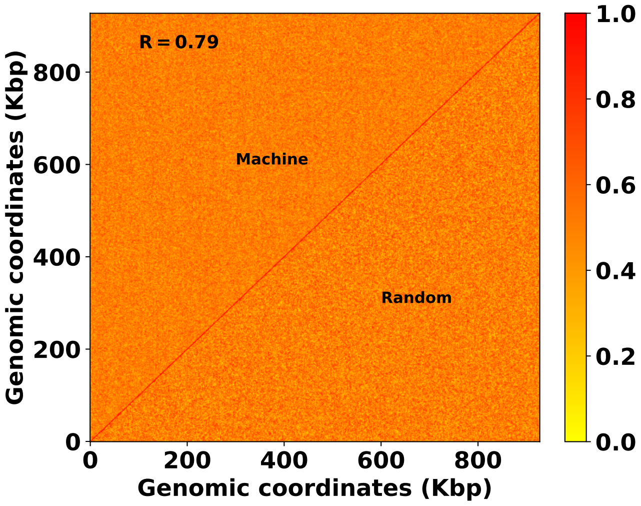


Figure S6: Comparison between the actual random matrix and ML-derived matrix. Here the dimension of the latent space $L_d = 40$

### References

[1] Aristidis Likas, Nikos Vlassis, and Jakob J Verbeek. The global k-means clustering algorithm. *Pattern recognition*, 36(2):451–461, 2003.

[2] Trupti M Kodinariya, Prashant R Makwana, et al. Review on determining number of cluster in k-means clustering. *International Journal*, 1(6):90–95, 2013.
